# Combining protein sequences and structures with transformers and equivariant graph neural networks to predict protein function

**DOI:** 10.1101/2023.01.17.524477

**Authors:** Frimpong Boadu, Hongyuan Cao, Jianlin Cheng

**Affiliations:** Department of Electrical Engineering and Computer Science, University of Missouri, Columbia, MO 65211, USA; Department of Statistics, Florida State University, Tallahassee, FL 32306, USA

## Abstract

**Motivation:** Millions of protein sequences have been generated by numerous genome and transcriptome sequencing projects. However, experimentally determining the function of the proteins is still a time consuming, low-throughput, and expensive process, leading to a large protein sequence-function gap. Therefore, it is important to develop computational methods to accurately predict protein function to fill the gap. Even though many methods have been developed to use protein sequences as input to predict function, much fewer methods leverage protein structures in protein function prediction because there was lack of accurate protein structures for most proteins until recently.

**Results:** We developed TransFun - a method using a transformer-based protein language model and 3D-equivariant graph neural networks to distill information from both protein sequences and structures to predict protein function. It extracts feature embeddings from protein sequences using a pre-trained protein language model (ESM) via transfer learning and combines them with 3D structures of proteins predicted by AlphaFold2 through equivariant graph neural networks. Benchmarked on the CAFA3 test dataset and a new test dataset, TransFun outperforms several state-of-the-art methods, indicating the language model and 3D-equivariant graph neural networks are effective methods to leverage protein sequences and structures to improve protein function prediction. Combining TransFun predictions and sequence similarity-based predictions can further increase prediction accuracy.

**Availability:** The source code of TransFun is available at https://github.com/jianlin-cheng/TransFun

## 1 Introduction

Proteins are essential macromolecules that carry out critical functions such as catalyzing chemical reactions, regulating gene expression, and passing molecular signals in living systems. It is critical to elucidate the function of proteins. However, even though various next-generation genome and transcriptome sequencing projects have generated millions of protein sequences, the experimental determination of protein function is still a low-throughput, expensive and time-consuming process. Thus, there is a huge gap between the number of proteins with known sequence and the number of proteins with known function, and this gap keeps increasing. As a result, it is important to develop computational methods to accurately predict the function of proteins.

Given the sequence of a protein and/or other information as input, protein function prediction methods aim to assign the protein to one or more function terms defined by Gene Ontology (GO)(Huntley et al., 2015). GO organizes function terms into three ontology categories: Biological Process (BP), Molecular Function (MF) and Cellular Component (CC). The terms in each of these ontology categories can be represented as a directed acyclic graph, in which parent nodes denoting broader (more general) function terms point to child nodes denoting more specific function terms.

Many protein function prediction methods use sequence or structure similarity to predict function, assuming proteins with similar sequences and structures likely have similar function. For example, GOtcha, Blast2GO (Conesa & Götz, 2008; Martin et al., 2004), PDCN (Wang et al., 2013) and DIAMONDScore use sequence alignment methods such as BLAST(Altschul et al., 1997) to search for homologous sequences with known function for a target protein and then transfer their known function to the target. COFACTOR and ProFunc (Laskowski et al., 2005; Zhang et al., 2017) use structure alignment to search for function-annotated proteins whose structures are similar to the target protein to transfer the function annotation. There are also some methods leveraging interactions between proteins or co-expression between genes to predict function, assuming that the proteins that interact or whose genes have similar expression patterns may have similar function. For instance, NetGO (You et al., 2019) transfers to a target protein the known function of the proteins that interact with it. All these nearest neighbor-based methods depend on finding related function-annotated proteins (or called templates) according to sequence similarity, structure similarity, gene expression similarity, or protein-protein interaction, which are often not available. Therefore, they cannot generally achieve high-accuracy protein function prediction for most proteins.

To improve the generalization capability of protein function prediction, advanced machine learning-based methods such as FFPred and labeler (Cozzetto et al., 2016; You et al., 2018) have been developed to directly predict the function of a protein from its sequence. However, most of these methods use hand-crafted features extracted from protein sequences to make prediction. Recently, several deep learning methods such as DeepGO (Kulmanov & Hoehndorf, 2020), DeepGOCNN (You et al., 2021), TALE(Cao & Shen, 2021), and DeepFRI (Gligorijević et al., 2021) were developed to predict protein function, leveraging deep learning’s capability to automatically extract features from input data. For instance, DeepFRI (Gligorijević et al., 2021) predicts the functions of proteins with a graph convolutional network by leveraging sequence features extracted by a long, short-term memory-based protein language model and structural features extracted from protein structures. However, DeepFRI uses either true protein structures from the Protein Data Bank(Berman et al., 2000) or homology-based structural models built by SWISS-MODEL as structure input. Because only a small portion of proteins have true structures or high-quality homology-based structural models, the method cannot be applied to most proteins. As AlphaFold2(Jumper et al., 2021; Varadi et al., 2022) can predict high-accuracy structures for most proteins, it is time to leverage AlphaFold2 predicted protein structures to advance protein function prediction.

In this work, we develop a method to use a pre-trained protein language transformer model to create embeddings from protein sequences and combine them with a graph representation constructed from AlphaFold predicted 3*D* structures through equivariant graph neural networks (EGNN) to predict protein function. We leverage the ESM language model(Elnaggar et al., 2021; Rao et al., 2021; Rives et al., 2019, 2021) trained on millions of protein sequences to generate good feature representations for protein sequences. The equivariant graph neural networks can capture the essential features of protein structures that are invariant to the rotation and translation of 3D protein structures to improve protein function. Our experiment shows that combining protein sequences and structures via the language transformer model and EGNN outperforms several state-of-the-art methods.

## 2 Methods

### 2.1 Datasets

We collected protein sequences with function annotations from the UniProt/Swiss-Prot database, released by February 23, 2022, amounting to a total of 566,996 proteins. We gathered their functional annotations from UniProt and the ontology graph data from the Open Biological and Biomedical Ontology (OBO) Foundry data repository. We also collected predicted structures of 542,380 proteins from the AlphaFold Protein Structure Database (AlphaFoldDB) published on January 12, 2022. To ensure consistency between the predicted structures from AlphaFoldDB and the corresponding proteins from UniProt, we compare their sequences and UniProt ID. All but 301 proteins have the same sequence. For the ones with different sequences, they usually only differ in a few residues. To make them consistent, we use the sequences extracted from the predicted structures as the final sequences.

The protein function annotations are described in the Gene Ontology (GO) terms. GO uses directed acyclic graphs (DAGs) to model the relationship between GO terms. The nodes represent the GO terms, and the links represent the relationship between the terms. GO provides three separate directed acyclic graphs (DAG) for each of the three ontologies (Biological Process (BP), Cellular Component (CC) and Molecular Function (MF)). For each protein, the specific GO terms provided in the UniProt function annotation file were first gathered. Then, their parent and ancestor terms in the GO DAG were also collected. The terms with the evidence codes (EXP, IDA, IPI, IMP, IGI, IEP, TAS, IC, HTP, HDA, HMP, HGI, HEP) were used as the function label for the protein according to the standard used in the Critical Assessment of Protein Function Annotation (CAFA)(Zhou et al., 2019).

The entire dataset above was filtered to retain only proteins with sequence length between 100 and 1022. We use a maximum length of 1022, because the pre-trained ESM model used in generating sequence embeddings can accept a protein sequence with the maximum length of 1022 residues. To avoid rare GO terms, we use only GO terms that have at least 60 proteins for training and test.

To compare our method with existing methods, we use the CAFA3 (Zhou et al., 2019) dataset as the independent test dataset because many methods have been tested on it. We removed all 3328 CAFA3 test proteins from our curated dataset and removed any protein in the dataset that has >=50% sequence identity with any protein in the CAFA3 test dataset. After the filtering, the curated dataset was used to train and validate TransFun. The trained method was then blindly tested on the test datasets.

We collected the predicted structures for the proteins in the CAFA3 test dataset (*CAFA3_test_dataset*) from AlphaFoldDB in the same way as for our curated dataset. If no predicted structure was found for a protein, we used AlphaFold2 to predict its structure. During the input feature generation, for a protein sequence with length > 1022 in the CAFA3 benchmark dataset, we divided it into smaller chunks of 1022 residues except for the last chunk for the language model to generate sequence embeddings that were concatenated together as the sequence embeddings for the entire protein sequence.

To investigate how sequence identity influence the accuracy of protein function prediction, we used mm2seq(Steinegger & Söding, 2017) to cluster the proteins in our curated dataset at the sequence identity thresholds of 30%, 50%, 90%. **Table 1** reports the total number of proteins in each function category, the total number of GO terms, and the number of protein clusters at each identity threshold.

**Table 1.** The statistics of the curated protein function prediction dataset. The first 3 columns are the GO ontology category, the total number of proteins in each category and the number of GO terms in each category. The remaining 4 columns list the number of protein clusters at each sequence identity threshold (0.3, 0.5, 0.9, 0.95).

| Ontology | # Protein | # GO Terms | Sequence Identity Threshold |  |  |  |
| --- | --- | --- | --- | --- | --- | --- |
|  |  |  | 0.3 | 0.5 | 0.9 | 0.95 |
| MF | 35,507 | 600 | 14,667 | 19,512 | 26,876 | 28,067 |
| CC | 50,340 | 547 | 20,679 | 26,808 | 36,721 | 38,509 |
| BP | 50,320 | 3774 | 20,180 | 26,647 | 37,536 | 39,348 |

Our final curated dataset was divided into training and validation sets. We randomly selected 5000 proteins with GO terms in all three ontology categories for validation.

We also collected new proteins released between March 2022 and November 2022 in UniProt as our second test dataset (*new_test_dataset*). This dataset has 702, 705 and 1084 proteins in CC, MF and BP respectively.

Given a set of proteins *D*_*l*_ = {(*p*_1_, *O*_1_), (*p*_1_, *O*_2_), *…* (*p*_*n*_, *O*_*n*_)}, where *p*_*i*_ is the *i*_*th*_ protein and *O*_*i*_ is its true function annotation labels (i.e., a set of GO terms). Our task is to predict *O*_*i*_ as accurately as possible. The function annotations are represented hierarchically with a general root term at the top. If a GO term *x* is associated with protein *p*_*i*_, then all the ancestor terms of *x* in the GO graph are also associated with protein *p*_*i*_. Therefore, the goal is to predict the sub-graph *g* in the GO graph *G* consisting of all the GO terms associated with the protein (Clark & Radivojac, 2013).

### 2.2 Protein Function Prediction Pipeline

We formulate the protein function task as a multi-label classification problem, where each protein may be assigned to one or more labels (GO terms). TransFun takes as input the sequence and predicted 3D structure of a protein and predicts the probability of GO terms for it in each GO category (**Figure 1**). TransFun consists of three main stages: **(1)** extracting a protein graph from a predicted structure (PDB), **(2)** generating the embeddings from a protein sequence, and **(3)** using a deep learning model to predict protein functions from the input data, which are described in Sections 2.3, 2.4, and 2.5.

**Figure 1.**
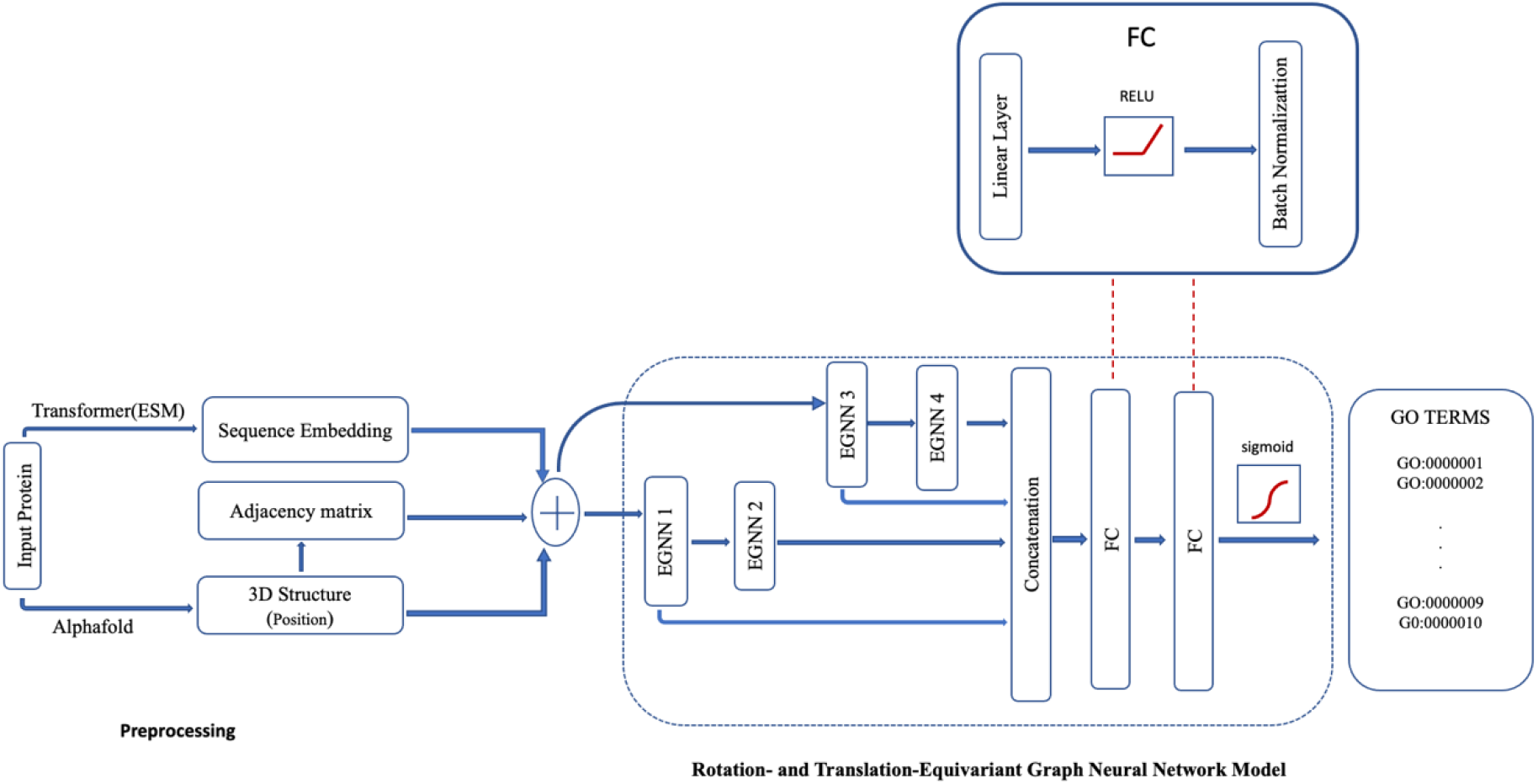
The protein function prediction pipeline of TransFun. The pipeline is divided into two main components, feature preprocessing(left) and neural network model (right). The input is a protein sequence. The output is the predicted probability of the GO terms for the protein.

### 2.3 Protein Graph Extraction from predicted structure

We construct a graph from the structure of a protein under consideration, represented as a *n* x *n* adjacency matrix, where *n* is the number of residues in the protein (**Figure 2)**. The nodes in the graph represent residues of the protein. Two types of edges are constructed between residues, using a distance threshold and K-nearest neighbor (KNN) approach. Given a protein graph *G* = (*V, E, X*), where *V* = {*v*_1_, *v*_2_, … *v*_*n*_ } represents the vertex set and *E* is the set of edges. The first condition for adding an edge to connect two nodes (*u, v*) is the Euclidean distance between their carbon alpha atoms |*u* − *v*| < *ϕ*, where *ϕ* is the distance threshold. In this work, we tested 5 distance thresholds, 4Å, 6Å, 8Å, 10Å & 12Å and chose 10Å as our final distance threshold as it yielded the best result. The second condition is *v* ∈ *Nk*, where *Nk* is the K nearest neighbors of node *u*. We set K to 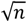 & 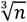, where n is the number of residues. Since both thresholds produce similar results, we use the latter to reduce computational cost. The graph constructed from a protein structure is stored in a binary adjacency matrix, where 0 means no edge and a 1 means there is an edge between two nodes. Self-loops (edges from a node to itself) are not allowed.

**Figure 2.**
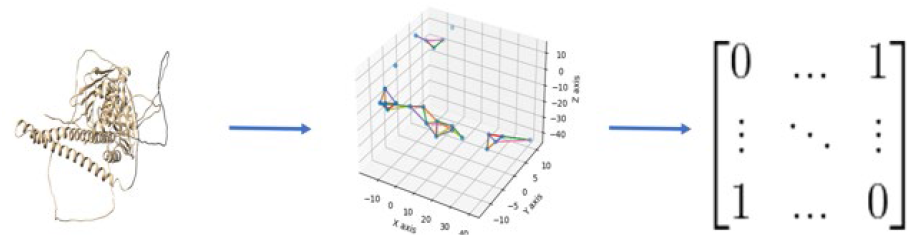
Constructing a graph from a protein structure. A graph is constructed from a protein structure using a distance threshold and K-nearest neighbor approach. The graph is stored in a binary adjacency matrix.

We also consider the situation when no protein structure is provided. In this case, a fully connected graph consisting of the edges between any two different nodes (residues) is constructed, which allows the deep model to infer appropriate edge weights during the training process, similar to the work (Kipf et al., 2018). However, this approach is computationally expensive, especially for large graphs.

### 2.4 Sequence Feature Extraction Using Transformer Language Model

We generate embeddings for the sequence of a protein using the ESM-1b(Rives et al., 2019, 2021) pre-trained protein language model. Per-residue embeddings are extracted for each residue (e.g., dimension: 21 × 1022) and per-sequence embedding for each whole sequence (e.g., dimension: 1022). The ESM-1b transformer takes as input the sequence of a protein and generates feature embeddings at several layers. We collect per-residue and per-sequence embeddings from the 33rd layer. The per-residue embedding for all the residues of a protein is an *R*^*l*×*d*^ tensor, where *l* is the sequence length and *d* is the embedding dimension. The per-sequence embedding is an aggregation over the per-residue embeddings and represents the features for the entire protein. We use the mean aggregator to compute the per-sequence embedding. ESM-1b was trained with the 1024 residue limit including start and end tokens (i.e., 1022 real residues without counting the start/end tokens). For a protein sequence with length > 1022, we divide the sequence into *n*/1022 chunks of length 1022 except for the last chunk that has a length of *n* % 1022, where *n* is the length of the sequence and generate an embedding for each chuck. The embeddings for all the chunks are concatenated as the embedding for the protein.

### 2.5 Rotation- and Translation-Equivariant Graph Neural Network (EGNN) Model

The deep graph neural network architecture of TransFun is composed of 4 blocks of rotation- and translation-equivariant graph neural networks (Satorras et al., 2021) (**Figure 1**), labeled as EGNN1, EGNN2, EGNN3, and EGNN4 respectively, each separated by a RELU activation function and Batch-normalization layer. Each EGNN block is made up of 4 equivariant graph neural network layers.

Each EGNN layer accepts a graph as input to update its features. The initial node features include the per-residue embeddings, and the (x, y, z) coordinates of each residue. We tested two optional features for the edges: (1) the distance between the two nodes of the edge; and (2) a binary number 0/1 indicating if the two residues are two adjacent residues connected by a peptide bond in the protein. However, the edge features do not improve the prediction accuracy on top of the node features and therefore are excluded in the final model of TransFun.

EGNN1 has an input dimension of 1022, equal to the feature embedding dimension for each node. It takes as input the protein graph with the per-residue embedding to generate a new embedding of dimension *C* and the refined coordinates of the nodes in the graph. *C* to set to the number of GO classes to be predicted. EGNN2 takes as input the protein graph with an output of dimension of *C* from EGNN1 as node embeddings and produces an output dimension of *C*/2. EGNN3 takes in as the initial per-sequence embedding of dimension 1022 for the protein to generate the new per-sequence embedding of dimension *C*/2. The last EGNN block (EGNN4) takes as input the output of dimension C/2 from EGNN3 to generate a per-sequence output of dimension of C/4.

The output embeddings (features) from EGNN1 and EGNN2 are aggregated by using a global mean pooling on the node features of each EGNN to obtain representative features for each protein. This is then concatenated with the per-sequence outputs of EGNN3 and EGNN4, resulting in a 2 *∗ C + C*/4 output features. The concatenated features are then passed through two fully connected (FC) linear layers, separated by batch normalization and RELU function to reduce the dimension to *C*. A sigmoid layer is used in the output layer to take the output of the last linear layer as input to predict the probability of each GO term.

### 2.6 Addressing Class Imbalanced Problems

The numbers of examples for different GO terms are very different. We use class weights to scale the training loss for GO terms appropriately to weigh less-represented GO terms (classes) more. The size of protein clusters in the training dataset is also imbalanced, where some clusters are very large, but some are very small. To reduce over the representation of proteins in a large cluster during training, we randomly sample one representative protein from each cluster for each training epoch. Although the representative protein sampled is similar in sequence to all the other proteins in the same cluster, their functional annotation may differ, especially when the sequence similarity is low. Therefore, we recompute the class weights per training epoch so that classes represented in the epoch are weighed appropriately.

### 2.7 Combining TransFun predictions with sequence similarity-based predictions

Several previous works (Cao & Shen, 2021; Kulmanov & Hoehndorf, 2020) combines an *ab initio* deep learning prediction method and a homology sequence similarity-based method such as DIAMONDScore (Buchfink et al., 2014, 2021) to improve protein function prediction. DIAMONDScore uses BLAST to search for homologous proteins and transfer their function annotations to a target protein under consideration. In this work, we also designed such a *composite* (or meta) method to combine the predictions from DIAMONDScore and the predictions of TransFun, which is called TransFun+. The score that TransFun+ predicts for a GO term is the weighted average of the score predicted by TransFun and the score predicted by DIAMONDScore. The weights were optimized on our curated validation dataset.

### 2.8 Evaluation Metrics

We use the two widely used metrics - Fmax and the Area Under the Precision-Recall curve (AUPR) - to evaluate the performance of our methods. The Fmax is the maximum F-measure computed over all the prediction thresholds. The F-measure for each threshold is computed as the harmonic mean of the precision (TP / (TP+FP)) and recall (TP / (TP + FN)), where TP is the number of true positives, FP the number of false positives, and FN the number of false negatives. The AUPR is computed by using the trapezoidal rule to approximate the region under the precision-recall curve.

## 3 Results and Discussions

After training and optimizing TransFun on our curated training and validation datasets, we blindly evaluated it on the new test dataset (new_test_dataset) and the CAFA3 test dataset (CAFA3_test_dataset) together with other methods.

### 3.1 Performance on CAFA3 test dataset

We compare TransFun with a Naïve method based on the frequency of GO terms, a sequence similarity-based method DIAMONDScore (Buchfink et al., 2014, 2021), and three recent deep learning methods DeepGO (Kulmanov & Hoehndorf, 2020), DeepGOCNN(Kulmanov & Hoehndorf, 2020), and TALE (Cao & Shen, 2021) on the CAFA3 test dataset in three function prediction categories (MF: molecular function; CC: cellular component, BP: biological process) in terms Fmax score and AUPR (**Table 2**). According to Fmax, TransFun performs best in MF and CC categories and second best in BP category. According to AUPR, TransFun performs best in MF and BP categories and second best in CC category. These results demonstrate that the sequence-based language transformer and 3D-equivariant graph neural network in TransFun can use protein sequence and structure together to improve function prediction over the existing methods.

**Table 2.** The results of TransFun and several other methods on the CAFA3 test dataset. TransFun was pretrained on the curated dataset whose proteins were clustered at sequence identity threshold of 50%. Bold numbers denote the best results.

| Method | Fmax |  |  | AUPR |  |  |
| --- | --- | --- | --- | --- | --- | --- |
|  | MF | CC | BP | MF | CC | BP |
| Naive | 0.295 | 0.539 | 0.315 | 0.138 | 0.373 | 0.197 |
| DIAMONDScore | 0.532 | 0.523 | 0.382 | 0.461 | 0.5 | 0.304 |
| DeepGO | 0.392 | 0.502 | 0.362 | 0.312 | 0.446 | 0.213 |
| DeepGOCNN | 0.411 | 0.582 | 0.388 | 0.402 | 0.523 | 0.213 |
| TALE | 0.548 | 0.654 | **0.398** | 0.485 | **0.649** | 0.258 |
| TransFun | **0.551** | **0.659** | 0.395 | **0.489** | 0.634 | **0.333** |

### 3.2 Impact of sequence identity on functional annotation

We compare the performance of TransFun on our curated validation datasets created using sequence identity thresholds of 30%, 50%, 90% respectively. The results are reported in **Table 3**. There is a slight increase of Fmax and AUPR when the sequence identify threshold is increased from 30% to 50% for molecular function (MF) and cellular component (CC), while the Fmax and AUPR for BP slightly decreases. This change may be also partially due to the difference in the test datasets at the two different sequence identity thresholds. However, the largely consistent results show that TransFun can work well when the sequence identity between the test protein and the training proteins is <= 30%. When the sequence identity threshold is increased from 50% to 90%, the performance is very similar, indicating when the sequence identity is higher than 50%, further increase sequence identity may not have a significant impact on the prediction accuracy.

**Table 3.** The results of TransFun on the test datasets having different identity thresholds with respect to the training data.

| Score | 30% |  |  | 50% |  |  | 90% |  |  |
| --- | --- | --- | --- | --- | --- | --- | --- | --- | --- |
|  | MF | CC | BP | MF | CC | BP | MF | CC | BP |
| Fmax | 0.509 | 0.619 | 0.394 | 0.53 | 0.631 | 0.37 | 0.53 | 0.606 | 0.367 |
| AUPR | 0.461 | 0.599 | 0.333 | 0.489 | 0.614 | 0.327 | 0.487 | 0.61 | 0.3 |

### 3.3 Performance on the new test dataset

**Table 4** reports the results of TransFun, Naïve, DIAMONDScore, two recent deep learning methods - DeepGOCNN and TALE, and three *composite* (meta) methods - DIAMONDScore – DeepGOPlus, TALE+ and TransFun+ on the new test dataset in the three ontology categories (MF, BP & CC) in terms of the Fmax score and AUPR score. Naïve, DIAMONDScore, DeepGOCNN, TALE, and TransFun are individual methods. DeepGOPlus, TALE+ and TransFun+ are composite (or meta) methods that combine the predictions of two individual methods (i.e., DeepGO + DIAMONDScore, TALE + DIAMONDScore, and TransFun + DIAMONDScore).

**Table 4.** The results on the new test dataset. Naïve, Diamond, DeepGOCNN, TALE and TransFun (green) are individual methods. DeepGOPlus, TALE+ and TransFun+ (blue) are composite methods. The best results of among the individual methods or among the composite methods are bold.

| Method | Fmax |  |  | AUPR |  |  |
| --- | --- | --- | --- | --- | --- | --- |
|  | CC | MF | BP | CC | MF | BP |
| Naïve | 0.560 | 0.275 | 0.283 | 0.404 | 0.135 | 0.173 |
| Diamond | 0.473 | 0.564 | 0.392 | 0.089 | 0.115 | 0.080 |
| DeepGOCNN | 0.595 | 0.440 | 0.361 | 0.545 | 0.307 | 0.240 |
| TALE | 0.607 | 0.512 | 0.344 | **0.613** | 0.480 | 0.257 |
| TransFun | **0.628** | **0.608** | **0.413** | 0.603 | **0.569** | **0.366** |
| DeepGOPlus | 0.623 | 0.635 | **0.460** | 0.562 | 0.549 | 0.339 |
| TALE+ | 0.619 | 0.635 | 0.431 | **0.633** | 0.613 | 0.344 |
| TransFun+ | **0.628** | **0.638** | 0.452 | 0.627 | **0.638** | **0.410** |

Among the four individual methods (Naïve, DIAMONDScore, DeepGOCNN, TALE, and TransFun), TrasnFun has the highest Fmax score of 0.628, 0.608, and 0.413 for CC, MF and BP, the highest AUPR score of 0.569 and 0.366 for MF and BP, and the second highest AUPR score of 0.603 for CC. TALE has the highest AUPR score of 0.621 for CC.

The three composite methods (DeepGOPlus, TALE+ and TransFun+) generally performs better than their individual counterpart (DeepGo, TALE and TransFun) in all the function categories in terms of both Fmax and AUPR except that TransFun and TransFun+ has the same Fmax score (i.e., 0.628) for CC. This indicates that combining the deep learning methods and sequence-similarity based methods can improve prediction accuracy. Among the three composite methods, TransFun+ performs best for CC and MF in terms of Fmax and for MF and BP in terms of AUPR, while DeepGOPlus performs best for BP in terms of Fmax and TALE+ performs best for CC in terms of AUPR.

The precision-recall curves of these methods on the new test dataset are plotted in **Figure 3**. It is worth noting that the deep learning methods such as TransFun, TALE and DeeoGOCNN perform much better than the sequence similarity-based method – DIAMONDScore, particularly in terms of AUPR. One reason is that DIAMONDScore has a much shorter precision-recall curve spanning a smaller range of recall values compared to the deep learning methods (see **Figure 3** for details).

**Figure 3.**
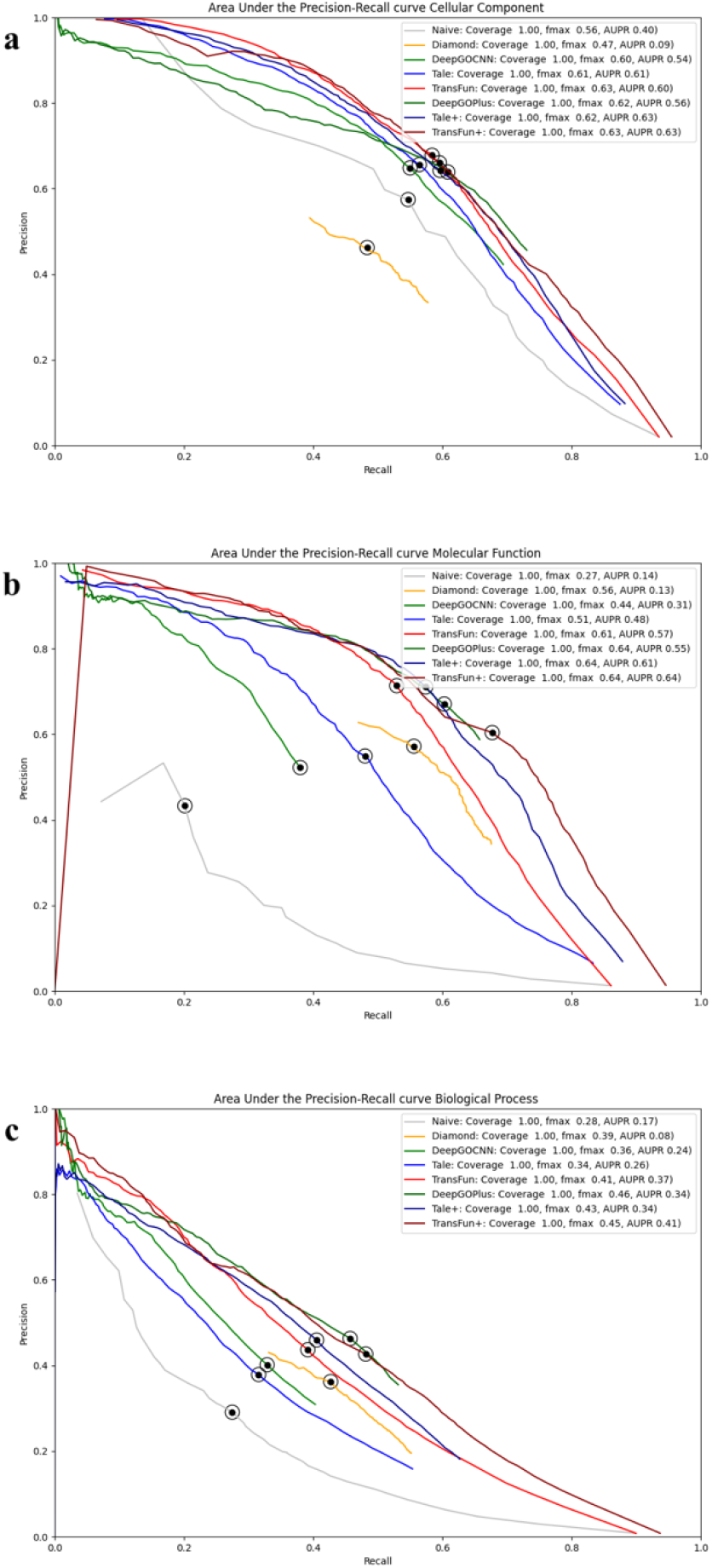
The precision-recall curves of the 8 methods on the new test dataset. The dot on the curves indicates where the maximum F score is achieved. The coverage is the percent proteins that a method makes predictions for.

### 3.4 Performance on human and mouse proteins

We compare the performance of TransFun, TransFun+ and the other methods on human proteins (**Table 5**) and mouse proteins (**Table 6**) in the new test dataset. In the dataset, there are 70, 35 and 34 human proteins in CC, MF and BP respectively, and there are 132, 87 and 158 mouse proteins for CC, MF and BP respectively.

**Table 5.** The results of the eight methods on human proteins in the new test dataset. Green denotes the individual methods and blue the composite methods. The best results in each type of methods are highlighted bold.

| Method | Fmax |  |  | AUPR |  |  |
| --- | --- | --- | --- | --- | --- | --- |
|  | CC | MF | BP | CC | MF | BP |
| Naive | 0.620 | 0.292 | 0.28 | 0.538 | 0.135 | 0.163 |
| Diamond | 0.509 | 0.516 | 0.445 | 0.085 | 0.087 | 0.055 |
| DeepGOCNN | 0.648 | 0.419 | 0.363 | 0.636 | 0.253 | 0.245 |
| TALE | 0.675 | 0.406 | 0.367 | **0.714** | 0.324 | 0.279 |
| TransFun | **0.686** | **0.538** | **0.468** | 0.694 | **0.471** | **0.445** |
| DeepGOPlus | 0.657 | 0.554 | 0.523 | 0.631 | 0.417 | 0.366 |
| TALE+ | **0.689** | 0.569 | 0.497 | **0.724** | 0.539 | 0.415 |
| TransFun+ | 0.684 | **0.612** | **0.553** | 0.719 | **0.557** | **0.499** |

**Table 6.** The results of the eight methods on mouse proteins in the new test dataset. Green denotes the individual methods and blue the composite methods. The best results in each type of methods are highlighted bold.

| Method | Fmax |  |  | AUPR |  |  |
| --- | --- | --- | --- | --- | --- | --- |
|  | CC | MF | BP | CC | MF | BP |
| Naive | 0.503 | 0.235 | 0.280 | 0.333 | 0.100 | 0.163 |
| Diamond | 0.471 | 0.569 | 0.379 | 0.087 | 0.119 | 0.082 |
| DeepGOCNN | 0.522 | 0.430 | 0.333 | 0.434 | 0.272 | 0.195 |
| TALE | 0.519 | 0.564 | 0.298 | 0.502 | 0.518 | 0.198 |
| TransFun | **0.558** | **0.576** | **0.355** | **0.517** | **0.532** | **0.289** |
| DeepGOPlus | **0.559** | 0.615 | **0.427** | 0.472 | 0.535 | 0.286 |
| TALE+ | 0.533 | **0.625** | 0.408 | 0.516 | 0.596 | 0.293 |
| TransFun+ | 0.557 | 0.624 | 0.403 | **0.529** | **0.618** | **0.352** |

The similar results are observed on the human and mouse proteins. Among the individual methods, TransFun performs better than the other methods in almost all function categories in terms of Fmax and AUPR.

The composite methods generally performs better than their corresponding individual methods. Among the three composite methods, TransFun+ performs best in most situations. These results are consistent with the results on all the proteins in the new test dataset (**Table 4**).

### 3.5 Performance on proteins longer than 1022 residues

Because TransFun and TransFun+ have to cut proteins longer than 1022 residues into pieces for the ESM-1b model to generate the sequence embedding features, we evaluated them and the other methods on the proteins longer than 1022 residues in the new test dataset. There are 49, 41 and 80 such proteins in CC, MF and BP respectively in the dataset. The results in **Table 7** show that TransFun yields the best performance in terms of AUPR for all three GO function categories among the individual methods and yields the best performance in terms of F_max_, for MF and BP. TransFun+ gives the best performance for CC and BP in terms of AUPR and the best performance for BP in terms of F_max_. DeepGOPlus gives the best results for CC in terms of F_max_, and TALE+ gives the best performance for MF in terms of F_max_. Compared with the results on all the proteins in **Table 4**, the performance of all the methods on the long proteins is generally lower than that on all the proteins with some exceptions, indicating that it is harder to predict the function of long proteins.

**Table 7.** The results on proteins longer than 1022 residues in the new test dataset. Green denotes the individual methods and blue the composite methods.

| Method | Fmax |  |  | AUPR |  |  |
| --- | --- | --- | --- | --- | --- | --- |
|  | CC | MF | BP | CC | MF | BP |
| Naive | 0.493 | 0.305 | 0.299 | 0.331 | 0.115 | 0.184 |
| Diamond | 0.560 | 0.583 | 0.427 | 0.060 | 0.093 | 0.086 |
| DeepGOCNN | **0.551** | 0.478 | 0.329 | 0.478 | 0.299 | 0.209 |
| TALE | 0.500 | 0.435 | 0.297 | 0.456 | 0.349 | 0.185 |
| TransFun | 0.550 | **0.556** | **0.399** | **0.525** | **0.492** | **0.353** |
| DeepGOPlus | **0.602** | 0.622 | 0.436 | 0.544 | 0.558 | 0.333 |
| TALE+ | 0.549 | **0.675** | 0.407 | 0.498 | **0.614** | 0.305 |
| TransFun+ | 0.564 | 0.593 | **0.443** | **0.563** | 0.610 | **0.396** |

### 3.6. A case study of protein function prediction

**Table 8** reports top 20 GO terms of biological process (BP) predicted for a protein (UniProt ID: A0A5S9MMK5) (length: 443; a putative transcription factor involved in morphogenesis) by four individual methods: Naïve, DeepGOCNN, TALE and TransFun. DIAMONDscore did not predict any result for this protein. All the top 20 GO terms predicted by TranFun are correct, while other methods made some incorrect predictions (red ones).

**Table 8.**
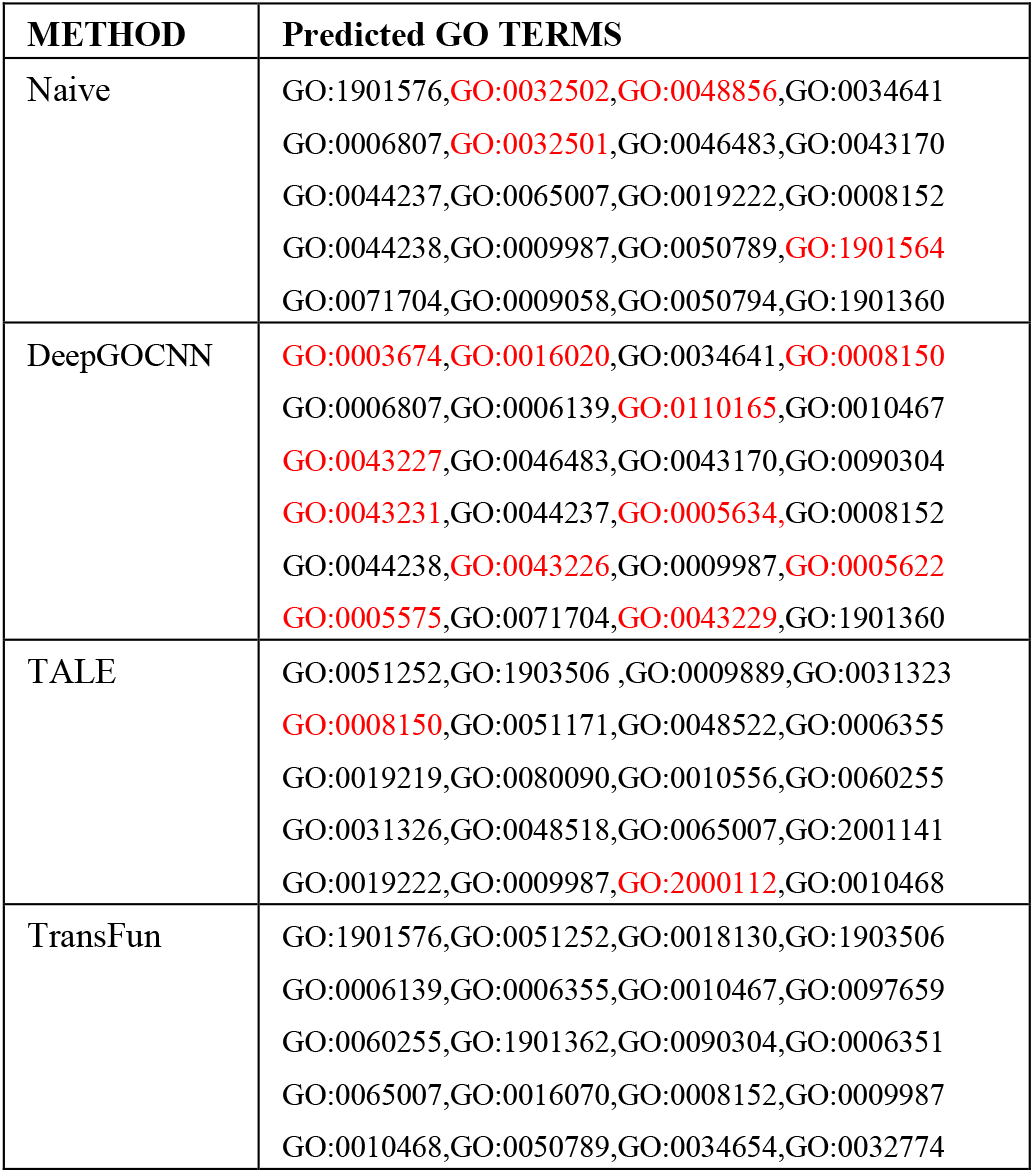
The GO terms for protein A0A5S9MMK5 by Naïve, DeepGOCNN, TALE, and TransFun. Black color denotes correct predictions, while red color denotes incorrect predictions.

## 4 Conclusion and Future Work

In this work, we developed TransFun for protein function prediction, using both protein structure and sequence information. TransFun uses transfer learning with a protein language model to extract sequence features and a graph representation to store structural features generated from AlphaFold predicted structures. The features are used by rotation/translation-equivariant graph neural networks to predict GO function terms for any protein. The method performs better than the sequence similarity-based and other deep learning methods on the two benchmark datasets. Moreover, TransFun can be combined with sequence similarity-based method to further improve prediction accuracy. In the future, we plan to use the multiple sequence alignment (MSA) of a target protein for the MSA-based language model (e.g., ESM-MSA) to generate extra embedding features for TransFun to see if they can further improve prediction accuracy. Another challenging issue facing protein function prediction is the lower prediction accuracy for more specific GO terms (the nodes at the lower levels of the gene ontology directed acyclic graph) because these terms have much fewer proteins associated with them than more general GO terms. More machine learning techniques and data preparation techniques are needed to address this imbalance problem because accurately predicting more specific GO terms is more useful for biological research than more general GO terms.

## Acknowledgements

We thank Elham Soltani Kazemi, Nabin Giri, Ashwin Dhakal, Raj Roy, Xiao Chen, and Alex Morehead for the assistance in preparing sequence data. The work was partly supported by the Department of Energy, USA (grant #: DE-AR0001213, DE-SC0020400, and DE-SC0021303),

National Science Foundation (grant #: DBI1759934 and IIS1763246), and National Institutes of Health (grant #.: R01GM093123 and R01GM146340).

